# fastDFE: fast and flexible inference of the distribution of fitness effects

**DOI:** 10.1101/2023.12.04.569837

**Authors:** Janek Sendrowski, Thomas Bataillon

## Abstract

Estimating the distribution of fitness effects (DFE) of new mutations is of fundamental importance in evolutionary biology, ecology, and conservation. However, existing methods for DFE estimation suffer from limitations, such as slow computation speed and limited scalability. To address these issues, we introduce fastDFE, a Python-based software package, offering fast and flexible DFE inference from site-frequency spectrum (SFS) data. Apart from providing efficient joint inference of multiple DFEs that share parameters, it offers the feature of introducing genomic covariates that influence the DFEs, and testing their significance. To further simplify usage, fastDFE is equipped with comprehensive VCF-to-SFS parsing utilities. These include options for site filtering and stratification, as well as site-degeneracy annotation and probabilistic 11 ancestral-allele inference. fastDFE thereby covers the entire workflow of DFE inference from the moment of acquiring a raw VCF file. Despite its Python foundation, fastDFE comprises a full R interface, including native R visualization capabilities. The package is comprehensively tested, and documented at fastdfe.readthedocs.io.

## 1 Introduction

The distribution of fitness effects (DFE) of new mutations is central to our understanding of how natural selection shapes genetic variation. Like its predecessors, fastDFE infers the DFE by contrasting the site-frequency spectrum (SFS) of putatively neutral (synonymous or intergenic) and selected (non-synonymous) sites (Keightley and Eyre-Walker, 2008; Galtier, 2016; Tataru et al., 2017). The intuition behind this approach is that beneficial alleles are more likely to segregate at higher frequencies than deleterious ones, implying that the SFS provides information on the fitness of new mutations. Summarizing all mutations’ fitness naturally gives rise to the distribution of fitness effects (DFE), which essentially is a probability distribution of the selection coefficients of mutations at selected sites.

To account for distortions to the selected SFS due to demography, the neutral SFS is commonly used, which is assumed to be subject to demography alone. Demography is either modeled explicitly, or by the introduction of nuisance parameters which are fit so as to re-scale the observed neutral SFS to match the standard Kingman SFS (Keightley and Eyre-Walker, 2008; Tataru et al., 2017). After accounting for demography, the remaining distortions to the selected SFS are assumed to be entirely due to selection.

SFS-based methods for estimating the DFE summarize population genetic variation by quantifying the number of alleles at specific frequencies, discarding details about the specific sites where these frequencies are exhibited. As a result, we obtain one DFE for all variants collectively. This approach provides limited information when considering a single DFE, but its utility significantly increases when comparing DFEs across multiple species, populations, or genomic regions (Moutinho et al., 2022; Chen et al., 2021; Latrille et al., 2023). Existing methods that can fit more than one SFS at a time are either notoriously slow and sometimes prone to numerical instabilities (Tataru and Bataillon, 2019), or offer only limited parameter sharing capabilities (Galtier, 2016). Here we introduce fastDFE which is specifically designed to facilitate and encourage the joint inference of multiple DFEs at once, allowing for the introduction of genomic covariates, and covering the entire workflow of DFE inference from the moment of acquisition of a raw VCF file.

## 2 Model & Enhancements

fastDFE primarily builds upon the model of polyDFE, overcoming its constraints such as extensive computational time and limited scalability (Tataru et al., 2017; Tataru and Bataillon, 2019). As such, it provides the same functionalities such as parametric bootstrapping, customizable DFE parametrizations, nested model comparison and joint inference. Similarly, the impact of demography is accounted for by introducing nuisance parameters (A.2).

To obtain the expected SFS given mutations having a specific selection coefficient, fastDFE makes use of the expected allele frequency sojourn times from Poisson random field (PRF) theory (Sawyer and Hartl, 1992; Sethupathy and Hannenhalli, 2008) (A.1). Let E [*P*_neut_(*i, S*)] denote the expected number of derived alleles at frequency *i* given (population-scaled) selection coefficient *S* (Figure A.1). Obtaining the expected SFS given a specific DFE requires integrating the expected SFS counts over the DFE, i.e., 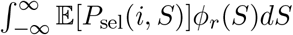, where *ϕ*_*r*_(*S*) denotes a density describing the DFE with regards to some chosen parametrization *r*. The actual result, i.e., inferring the DFE given an SFS is then achieved by maximum-likelihood optimization with respect to *r*, where a composite Poisson likelihood for the SFS is used (A.4).

In fastDFE, the above integral is discretized, precomputed and cached, leading to significant performance improvements over polyDFE while producing very similar results (Table B.1, Figures B.1 & B.2). One issue with polyDFE is its numerical integration procedure, which has difficulties to properly capture the DFE’s mass in regions where it is highly concentrated. One example of this is the oft-used G distribution with low shape parameters. fastDFE circumvents this problem by describing the DFE by means of its cumulative distribution function (CDF), thus providing more reliable estimates (A.5).

When inferring DFEs from multiple datasets, it is essential to standardize the derivation of the SFS to ensure results are directly comparable. To aid this process, fastDFE comes equipped with a VCF parser, enabling the extraction of the necessary SFS input data from raw VCF files. In order to obtain multiple SFS, a single VCF file can optionally be stratified with regards to some genomic properties, the stratification into a neutral and selected SFS being one example. To further aid the data preparation process, sites can be filtered and annotated with regards to degeneracy and ancestral state. Distinguishing between ancestral and derived alleles is important as it greatly improves the accuracy of DFE estimates, especially for beneficial mutations. The implemented ancestral allele annotation utility uses outgroup information, and follows the probabilistic model of EST-SFS (Keightley and Jackson (2018)). At last, utilities are provided to determine the number of mutational target sites when monomorphic sites are not present in the VCF file at hand.

## 3 Examples

### 3.1 Basic inference

In the following Python code snippet, we demonstrate how to use fastDFE to infer the DFE for a single population. Firstly, we define the spectra for both neutral (`sfs neut`) and selected (`sfs sel`) sites for a sample size of *n* = 8, leveraging data from a Scandinavian silver birch study (Sendrowski (2022)). We also parametrize the DFE using [GammaExpParametrization](https://fastdfe.readthedocs.io/en/latest/modules/parametrization.html#fastdfe.parametrization.GammaExpParametrization)—a common parametrization consisting of a mixture of a gamma and an exponential distribution for deleterious and beneficial selection coefficients, respectively. We use this information to instantiate a [BaseInference](https://fastdfe.readthedocs.io/en/latest/modules/inference.html#fastdfe.base_inference.BaseInference) object, which serves to infer a single DFE.

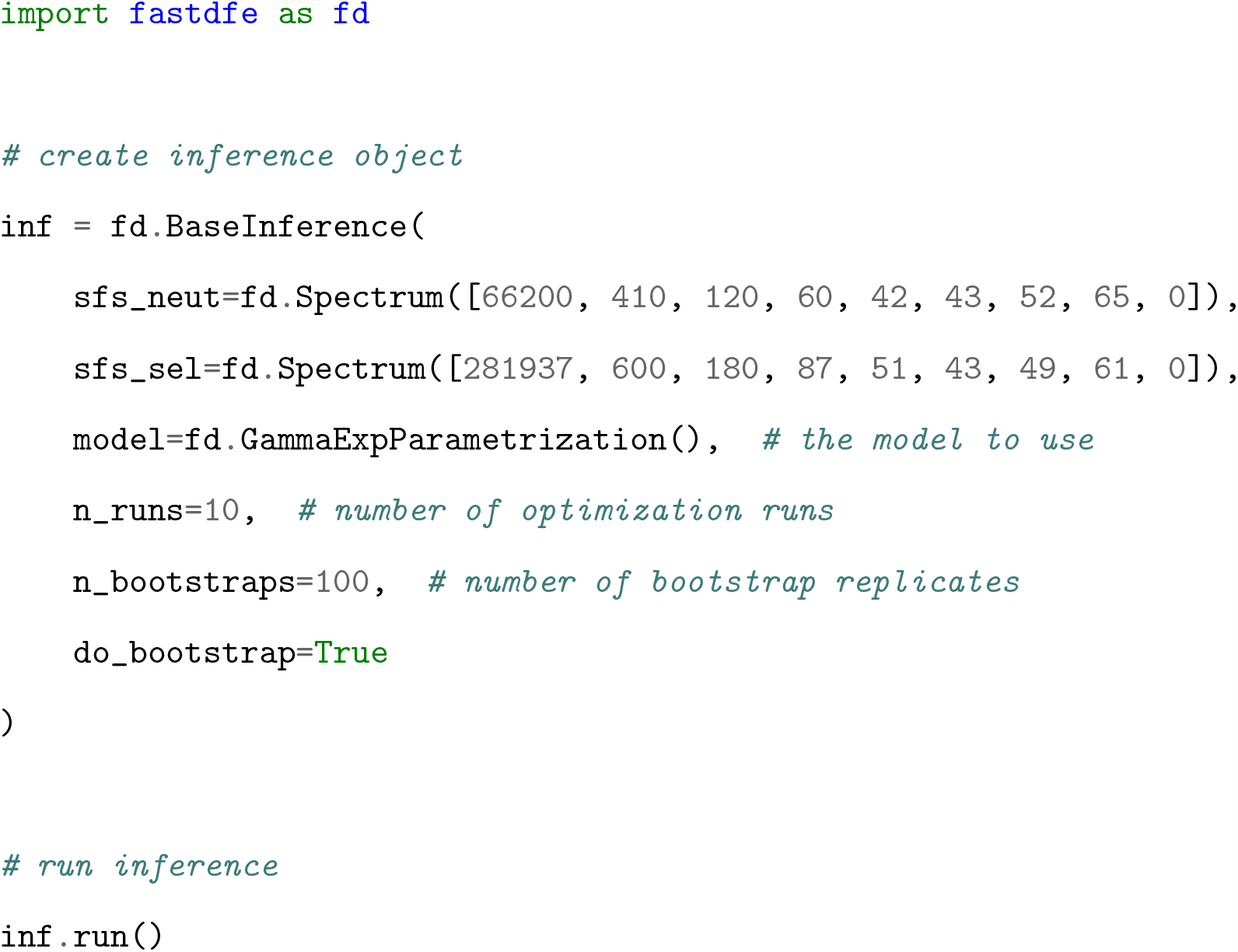

Upon carrying out the inference process, we proceed to visualize the results (Figure 1). Note that an equivalent R script would have significant similarities to the provided Python example.

**Figure 1:**
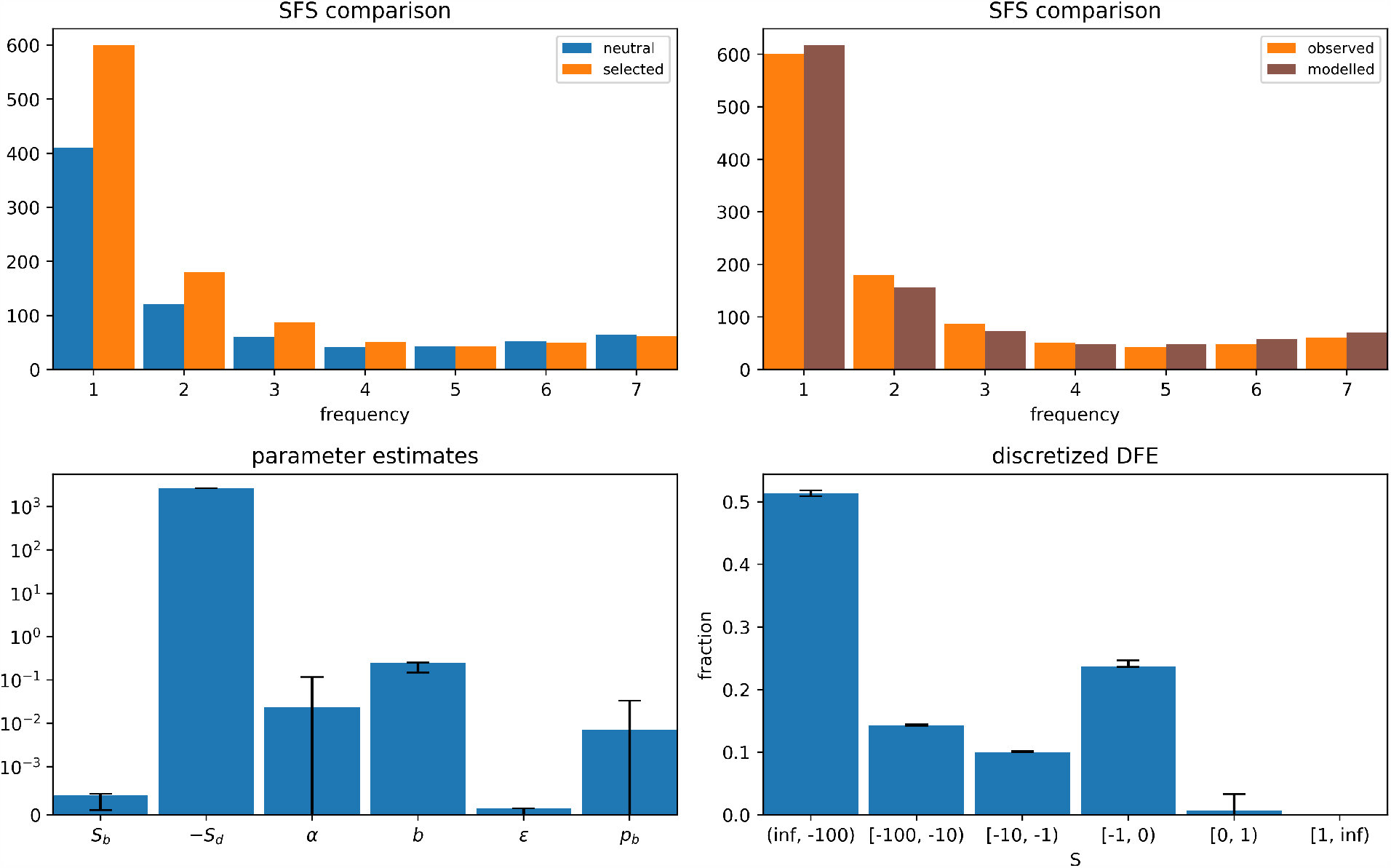
Output from code snippet in Section 3.1. **Top left**: observed neutral and selected SFS. **Top right**: observed (selected) and modelled (selected) SFS. **Bottom left**: parameter estimates. For `GammaExpParametrization`, *S*_*d*_ and *S*_*b*_ represent the average strength of deleterious and beneficial mutations, respectively. Parameter *b* denotes the shape of the gamma distribution and *p*_*b*_ the probability of a mutation being beneficial, whereas *ϵ* is the ancestral misidentification parameter (A.3). *α* denotes the expected proportion of beneficial non-synonymous substitutions, and is not a real parameter, but rather a function of the inferred DFE. **Bottom right**: discretized DFE estimate; The vertical bars indicate 95% confidence intervals.

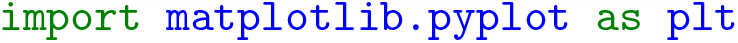
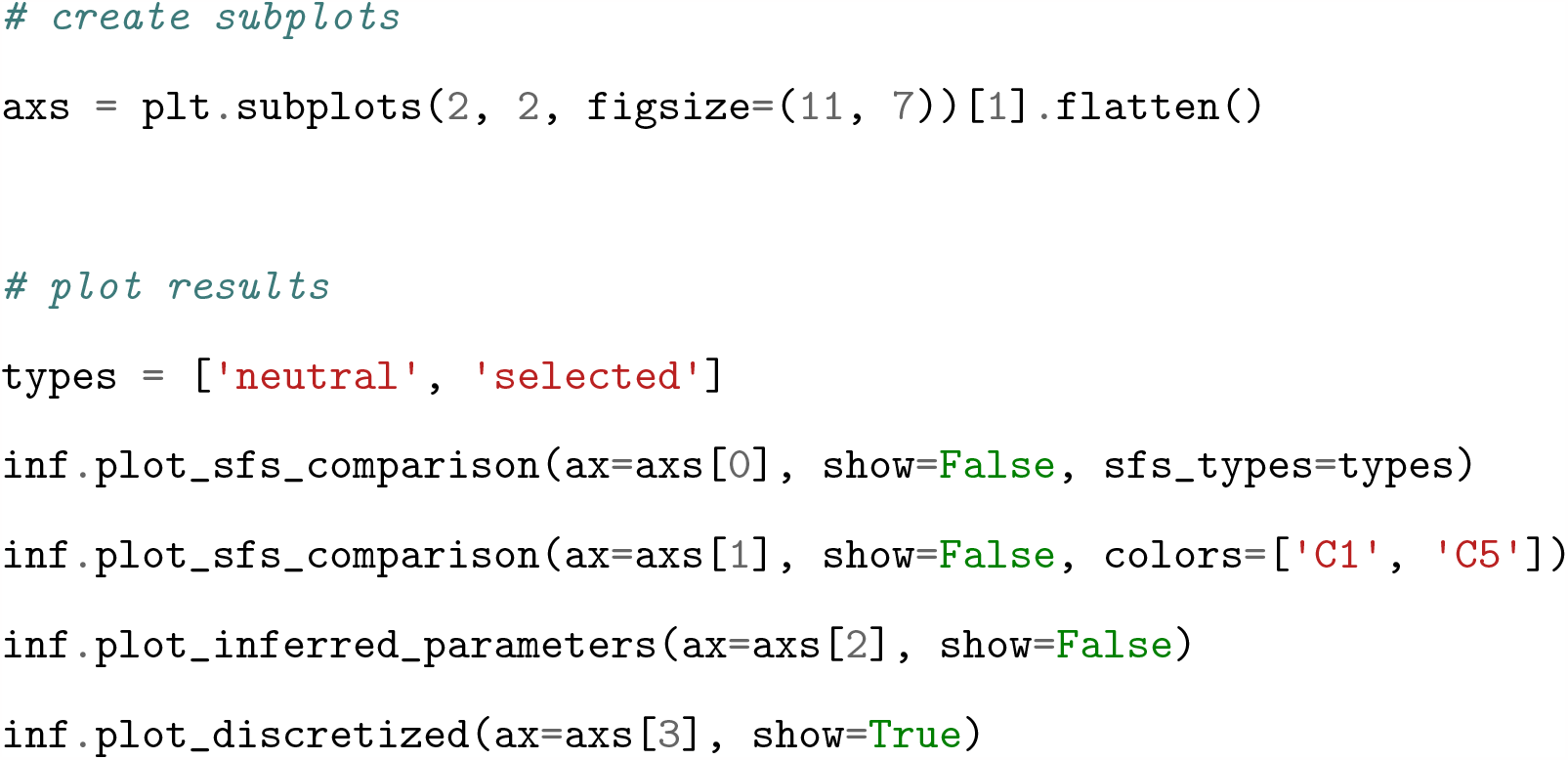

### 3.2 VCF parsing

fastDFE offers a versatile parser extracting SFS from VCF files while allowing for stratifications of the SFS. The stratified spectra can be directly used for either joint or marginal DFE estimation. In the following code example, we obtain the SFS used in Section 3.1. We determine the site-degeneracy on-the-fly by passing [DegeneracyAnnotation](https://fastdfe.readthedocs.io/en/latest/modules/annotation.html#fastdfe.annotation.DegeneracyAnnotation), which requires both a FASTA and a GFF file to be specified. [DegeneracyStratification](https://fastdfe.readthedocs.io/en/latest/modules/parser.html#fastdfe.parser.DegeneracyStratification) is then used to stratify the SFS into *selected* and *neutral* sites. Ancestral alleles are determined by passing [MaximumLikelihoodAncestralAnnotation](https://fastdfe.readthedocs.io/en/latest/modules/annotation.html#fastdfe.annotation.MaximumLikelihoodAncestralAnnotation). Since the specified VCF file only contains biallelic sites, we also provide [TargetSiteCounter](https://fastdfe.readthedocs.io/en/latest/modules/parser.html#fastdfe.parser.TargetSiteCounter), which determines the number of monomorphic sites for each stratification by sampling sites from the provided FASTA file. The resulting SFS are depicted in Figure C.1.

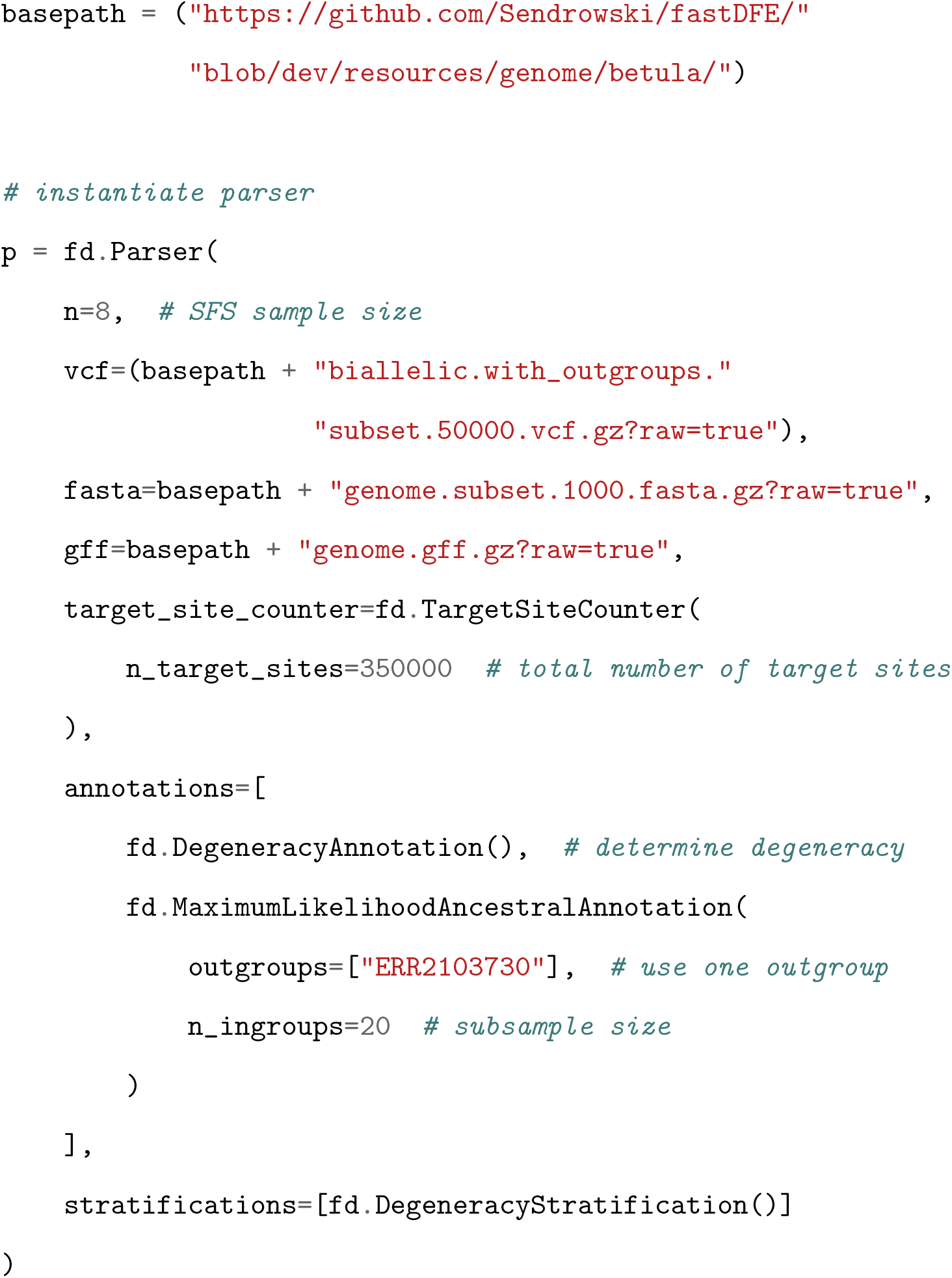
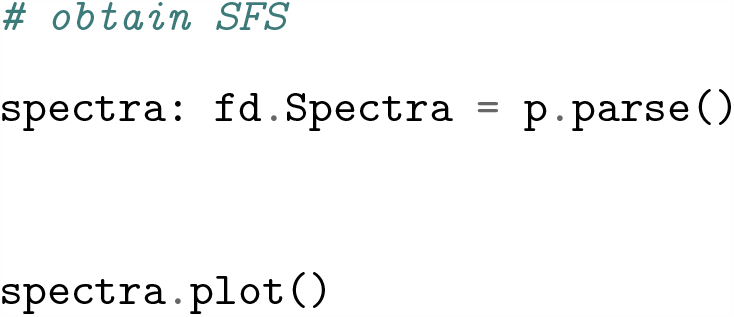

### 3.3 Joint inference

In comparative analyses, the ability to stratify and associate site-frequency spectra with covariates (such as genomic features or species-specific properties) can prove very useful. Joint inference enables the sharing of an arbitrary number of parameters across multiple datasets, henceforth referred to as types. Parameters that are not shared are estimated marginally, i.e., for each type separately. However, when using many types with marginal parameters, the large number of parameters to be optimized can make it difficult to find good optima. Covariates address this by introducing a customizable, typically linear, relationship with a specific parameter across all types. This reduces the number of additional parameters to one per covariate, and allows for the testing of covariate significance through a likelihood ratio test against the null model, where the parameter in question is constant across all types.

In the example below, we access a configuration file containing data from a study testing the adaptive walk model of gene evolution in *A. thaliana* (Moutinho et al., 2022). The underlying premise is that younger genes, due to their farther distance from their fitness optimum, are more likely to experience mutations with large fitness effects compared to older genes. The study provides site-frequency spectra stratified by gene, which are associated with a number of proxies for gene age. Notably, we focus on the relative solvent accessible (RSA) as our proxy of interest. The spectra for each gene, after being downsampled to 20 individuals, were grouped into 10 quantiles based on their RSA score. The covariate for each bin was taken to be the mean RSA score over all its genes. A linear relationship is then introduced with *S*_*d*_—the average strength of deleterious mutations (cf. this script). All remaining parameters are specified to be shared (i.e. constant) across all types. The significance of the included covariate is determined by comparing the likelihood against the null model in which *S*_*d*_ is constant across all bins (Figure 2).

**Figure 2:**
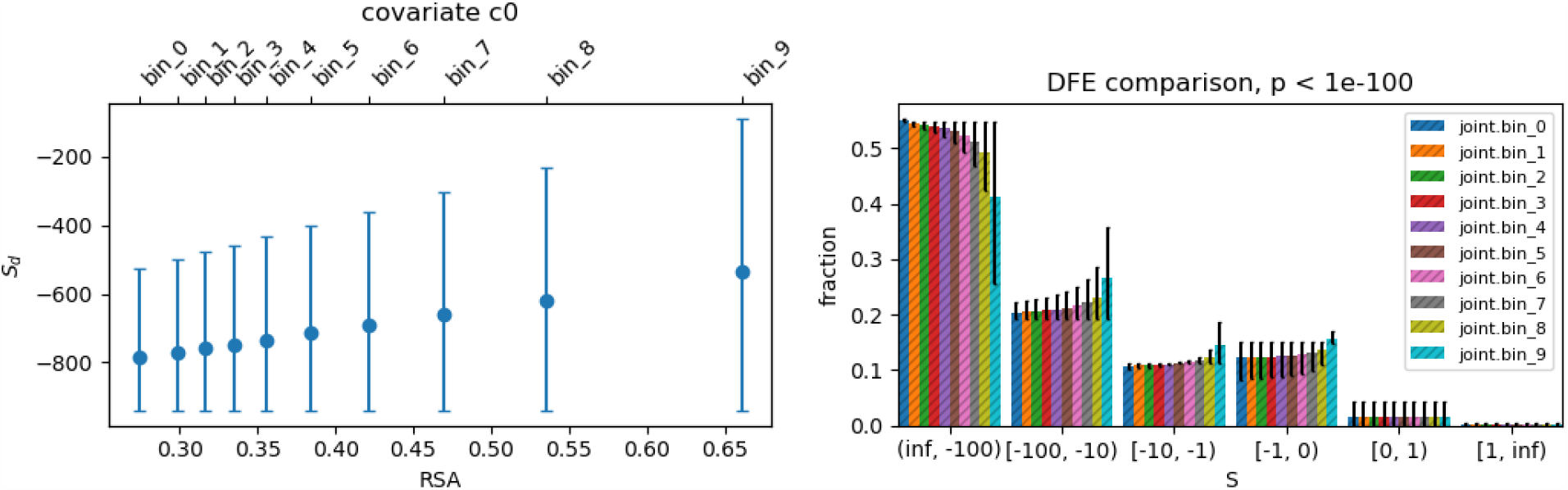
Output from code snippet in Section 3.3. **Left**: values of *S*_*d*_ vs. RSA score for the different bins. *S*_*d*_ is restricted to vary linearly with RSA. **Right**: inferred joint DFE. The low p-value (indicated in the plot title) confirms the high significance of covariate inclusion; Vertical bars indicate 95% confidence intervals.

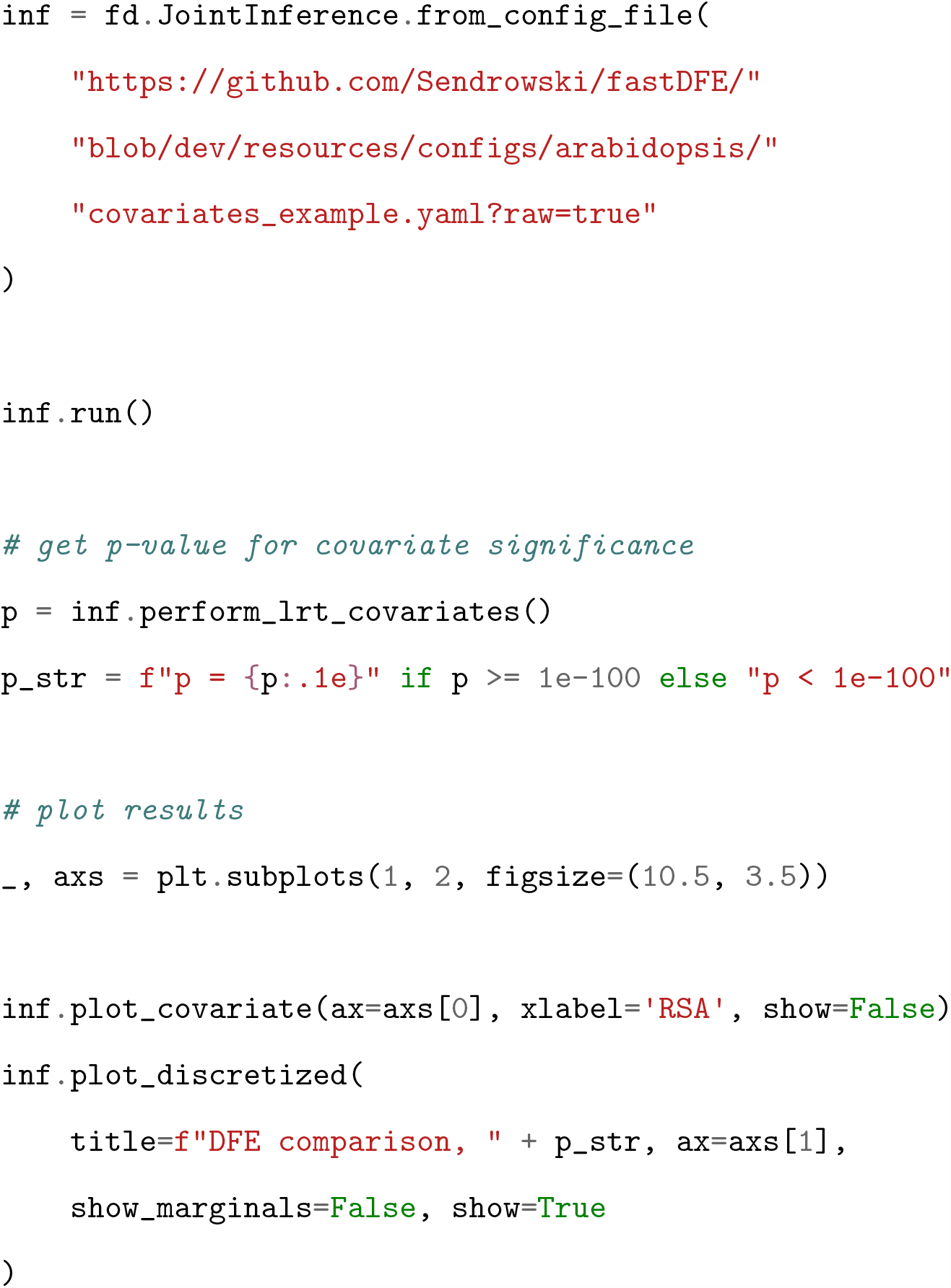

## 4 Discussion

fastDFE aims to improve on existing DFE estimation software, particularly in the realm of joint inference and data preparation. It offers a very efficient solution for joint inference—both performance-wise and by allowing for the introduction of covariates. It is thus well-suited for comparative analyses, whether across species, populations, or varying genomic contexts. The VCF parsing and annotation features greatly facilitate the data preparation process, potentially minimizing data handling errors while making different datasets more comparable. Customizable DFE parametrizations by means of the cumulative distribution function (CDF) also offer increased flexibility and reliability. The availability of a Python and R interface instead of a traditional command line approach aims to make DFE estimation more customizable. In conclusion, we hope that fastDFE will contribute to new insights in evolutionary biology, ecology, and conservation—especially within the scope of comparative studies. The software is freely available at github.com/Sendrowski/fastDFE, and comprehensive documentation as well as usage examples can be found at `fastdfe.readthedocs.io`.

## Acknowledgements

We would like to thank Sylvain Glémin for discussion on model implementation and Julien Dutheil for his helpful comments on the manuscript. This work has been supported by the Novo Nordisk Foundation (Data Science Collaborative Research Programme, grant 0069105). We also acknowledge Martin Lascoux and Luis Leal (Formas, grant 2020-01456) for kindly providing the birch dataset.

