## Appendix for "fastDFE: fast and flexible inference of the distribution of fitness effects"

### A Model and Implementation

#### A.1 Basic Model

As in polyDFE, the (expected) neutral ($P_{\text{neut}}(i)$) and selected ($P_{\text{sel}}(i)$) counts at derived allele frequency $i$ are modeled using Poisson random field theory:

$$\mathbb{E}[P_{\text{neut}}(i)] = \frac{\theta}{i}, \tag{1}$$

$$\mathbb{E}[P_{\text{sel}}(i)] = \theta \int_0^1 B(i,n,x)H(S,x)dx, \tag{2}$$

where $\theta = 4N_e\mu$ is the population-scaled mutation rate,

$$B(i,n,x) = \binom{n}{i} x^i (1-x)^{n-i}$$

is the binomial probability of observing $i$ derived alleles in a sample of size $n$ when the real allele frequency is $x$, and

$$H(S,x) = \frac{1-e^{-S(1-x)}}{x(1-x)(1-e^{-S})}$$

is the mean time a new semi-dominant mutation of population-scaled selection coefficient $S = 4N_e s$ spends between $x$ and $x + dx$ (Tataru et al., 2017). Equation 2 can be solved more efficiently by rewriting it as

$$\mathbb{E}[P_{\text{sel}}(i)] = \theta \frac{n}{i(n-i)} \frac{1 - e^{-S} \cdot 1F1(i,n,S)}{1-e^{-S}},$$

where $1F1$ denotes the confluent hypergeometric function of the first kind (Tataru et al., 2017). To circumvent numerical errors, we use approximations for $\mathbb{E}[P_{\text{sel}}(i)]$ at points where $S$ is either close to 0 or very negative (Figure A.1). Specifically, we use:

$$\mathbb{E}[P_{\text{sel}}(i)] \approx \frac{\theta}{i}, \qquad S \in (-\epsilon, \epsilon),$$

$$\mathbb{E}[P_{\text{sel}}(i)] \approx \theta \cdot \frac{n}{i(n-i)} \cdot 1F1(i,n,S), \qquad S \ll 0.$$

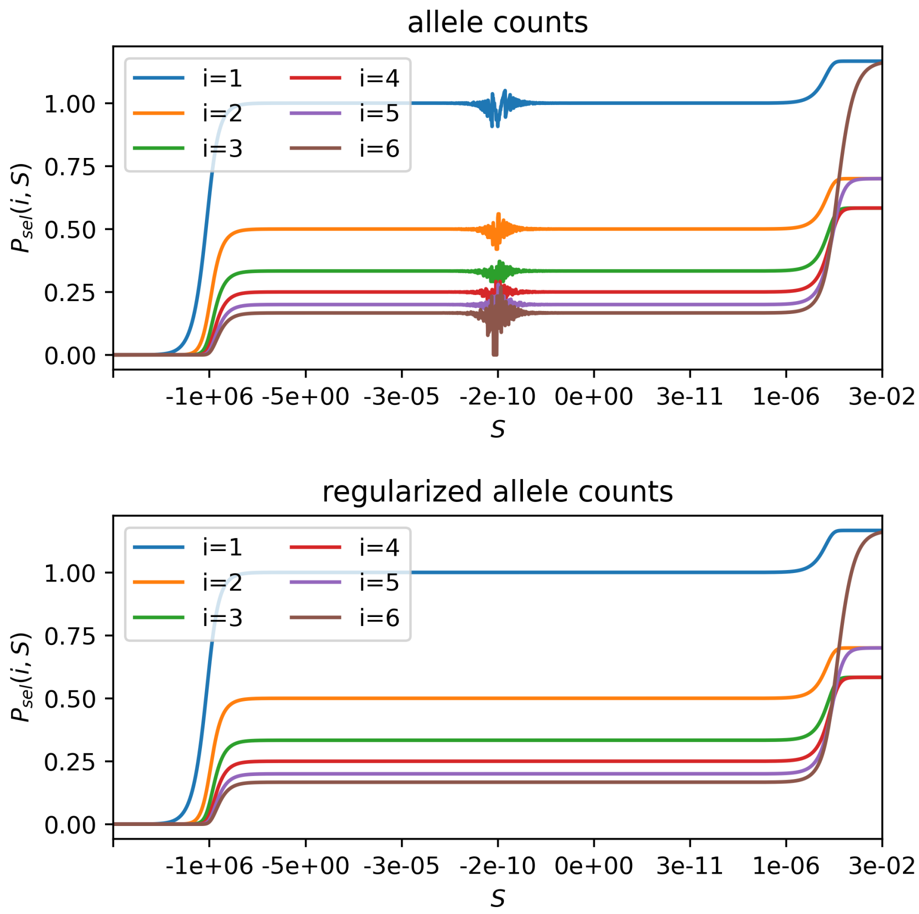


Figure A.1: Regularized and non-regularized expected allele counts at different derived allele counts $i$ for $\theta = 1$ and a sample size of $n = 7$, (Equation 2). The non-regularized version is subject to numerical instabilities.

##### A.2 Demographic Correction

We accommodate demographic effects by rescaling the observed selected SFS ($\hat{P}_{\text{sel}}$), using the ratio of the expected neutral counts ($P_{\text{neut}}$) (without demography) to the observed neutral counts $\hat{P}_{\text{neut}}$ (including demography). This method, while similar to the approach of polyDFE, solely uses the observed neutral SFS to correct for demography, resulting in faster computation while maintaining a similar level of accuracy provided that a reasonably large number of alleles counts is given. We have

$$\mathbb{E}[\hat{P}'_{\text{sel}}(i)] = \mathbb{E}[\hat{P}_{\text{sel}}(i)] \cdot \frac{\mathbb{E}[P_{\text{neut}}(i)]}{\mathbb{E}[\hat{P}_{\text{neut}}(i)]},$$

where $\mathbb{E}[\hat{P}'_{\text{sel}}(i)]$ are the demography-corrected observed selected counts at derived allele frequency $i$.

##### A.3 Ancestral Misidentification

As in polyDFE, the model can optionally incorporate ancestral misidentification. Let $\epsilon$ be the probability of mistakenly classifying alleles as derived when they are in fact ancestral, and vice versa. That is,

$$\mathbb{E}[\hat{P}'_z(i)] = (1-\epsilon)\mathbb{E}[P_z(i)] + \epsilon\mathbb{E}[P_z(n-1)],$$

for $z \in \{\text{sel}, \text{neut}\}$.

##### A.4 Likelihood

The selected modeled SFS is obtained by integrating over the DFE, i.e,

$$\mathbb{E}[P_{\text{sel}}(i)] = \int_{-\infty}^{\infty} \mathbb{E}[P_{\text{sel}}(i,S)]\phi_r(S)dS, \tag{3}$$

where $\phi_r(S)$ is a density describing the DFE with regards to some parametrization $r$. The parameters of $\phi_r(S)$ are then optimized numerically using the composite Poisson likelihood

$$\mathcal{L}(\theta) = \prod_{i=1}^{n-1} \text{Pois}(\hat{P}_{\text{sel}}(i), P'_{\text{sel}}(i)) \tag{4}$$

where $\mathrm{Pois}(k;\mu) = \frac{e^{-\mu}\mu^k}{k!}$ is the Poisson distribution, and $\hat{P}_{\mathrm{sel}}(i)$, and $P'_{\mathrm{sel}}(i)$ are the observed and modeled counts, respectively (Sawyer and Hartl, 1992; Sethupathy and Hannenhalli, 2008). For joint inference, we furthermore assume the likelihood of the component SFS to be independent.

#### A.5 Discretization

The expected allele counts $P_{\mathrm{sel}}(i, S)$ are rather well-behaved with regards to $S$ (Figure A.1), so that the integral in Equation 3 can be linearized. Furthermore, $\mathbb{E}[P_{\mathrm{sel}}(i, S)]$ does not change over the course of the optimization procedure, so that it can be precomputed and cached. The integral in Equation 3 can thus be discretized. Let $s_0, s_1, \ldots, s_k$ be the edges dividing the interval $(L, U)$ into $k$ bins, where $(L, U)$ is chosen such that it contains almost all of the mass of $\phi_r(S)$ for all $r$. We have

$$\begin{aligned}
\mathbb{E}[P_{\mathrm{sel}}(i)] &= \sum_{j=1}^{k} \mathbb{E}[P_{\mathrm{sel}}^{j}(i)] \\
&= \sum_{j=1}^{k} \int_{s_j}^{s_{j-1}} \mathbb{E}[P_{\mathrm{sel}}(i, S)]\phi(S)dS \\
&\approx \sum_{j=1}^{k} (s_{j-1} - s_j) \cdot \mathbb{E}\left[\frac{P_{\mathrm{sel}}(i, s_{j-1}) + P_{\mathrm{sel}}(i, s_j)}{2}\right] \cdot (\Phi_r(s_j) - \Phi_r(s_{j-1}))\,,
\end{aligned}$$

where $\Phi_r$ is the cumulative distribution function of the DFE. For an SFS sample size of $n$, we can then precompute the $(n \times k)$ matrix

$$P_{\mathrm{sel}} = \begin{bmatrix} P_{\mathrm{sel}}^{11} & P_{\mathrm{sel}}^{12} & \cdots & P_{\mathrm{sel}}^{1k} \\ P_{\mathrm{sel}}^{21} & P_{\mathrm{sel}}^{22} & \cdots & P_{\mathrm{sel}}^{2k} \\ \vdots & \vdots & \ddots & \vdots \\ P_{\mathrm{sel}}^{n1} & P_{\mathrm{sel}}^{n2} & \cdots & P_{\mathrm{sel}}^{nk}, \end{bmatrix} \tag{5}$$

where

$$P_{\mathrm{sel}}^{ij} = (s_{j-1} - s_j) \cdot \mathbb{E}\left[\frac{P_{\mathrm{sel}}(i, s_{j-1}) + P_{\mathrm{sel}}(i, s_j)}{2}\right]. \tag{6}$$

Now for each optimization iteration, we compute

$$w = \begin{bmatrix} w_1 \\ w_2 \\ \vdots \\ w_k \end{bmatrix} = \begin{bmatrix} \Phi_r(s_1) - \Phi_r(s_0) \\ \Phi_r(s_2) - \Phi_r(s_1) \\ \vdots \\ \Phi_r(s_k) - \Phi_r(s_{k-1}) \end{bmatrix}, \tag{7}$$

so that

$$P_{\text{sel}} w = \begin{bmatrix} w_1 P_{\text{sel}}^{11} & + & w_2 P_{\text{sel}}^{12} & + & \cdots & + & w_k P_{\text{sel}}^{1k} \\ w_1 P_{\text{sel}}^{21} & + & w_2 P_{\text{sel}}^{22} & + & \cdots & + & w_k P_{\text{sel}}^{2k} \\ \vdots & & \vdots & & \ddots & & \vdots \\ w_1 P_{\text{sel}}^{n1} & + & w_2 P_{\text{sel}}^{n2} & + & \cdots & + & w_k P_{\text{sel}}^{nk} \end{bmatrix} \tag{8}$$

is the resulting SFS. This greatly accelerates the numerical optimization as we are solely required to compute $w$ for each run. In practice, the edges $s_i$ are log-spaced for negative and positive $S$ separately to ensure a smaller spacing where the integral in Equation 3 varies more. This procedure yields very accurate estimates, and describing the DFE by means of its CDF ensures we can properly capture the mass, even for very leptokurtic or discontinuous distributions, which can prove problematic with conventional numerical integration methods.

### B Comparison with polyDFE

| dataset | full/del DFE | $\epsilon$ | fastDFE (s) | polyDFE (s) |
|---|---|---|---|---|
| *B. pendula* | full | ✓ | 26.69 | 145 638.58 |
| *B. pendula* | full | | 21.43 | 105 922.76 |
| *B. pendula* | deleterious | ✓ | 9.10 | 98 984.59 |
| *B. pendula* | deleterious | | 6.89 | 10 487.42 |
| *B. pubesens* | full | ✓ | 20.59 | 101 215.85 |
| example 1 | full | ✓ | 26.67 | 29 175.01 |
| example 2 | full | ✓ | 30.11 | 69 988.49 |
| example 3 | full | ✓ | 27.93 | 32 644.71 |

Table B.1: Comparison of unparallelized runtime in seconds for fastDFE and polyDFE across an array of datasets. A single core equipped with 8GB of memory was utilized. The runtime difference is substantial and often renders joint inference prohibitively expensive for polyDFE. *B. pendula* and *B. pubesens* constitute a dataset of Scandinavian silver birches (Sendrowski, 2022). Examples 1-3 can be found on the official GitHub repository for polyDFE (`github.com/paula-tataru/polyDFE/tree/master/input`). The column with $\epsilon$ indicates the inclusion of ancestral misidentification.

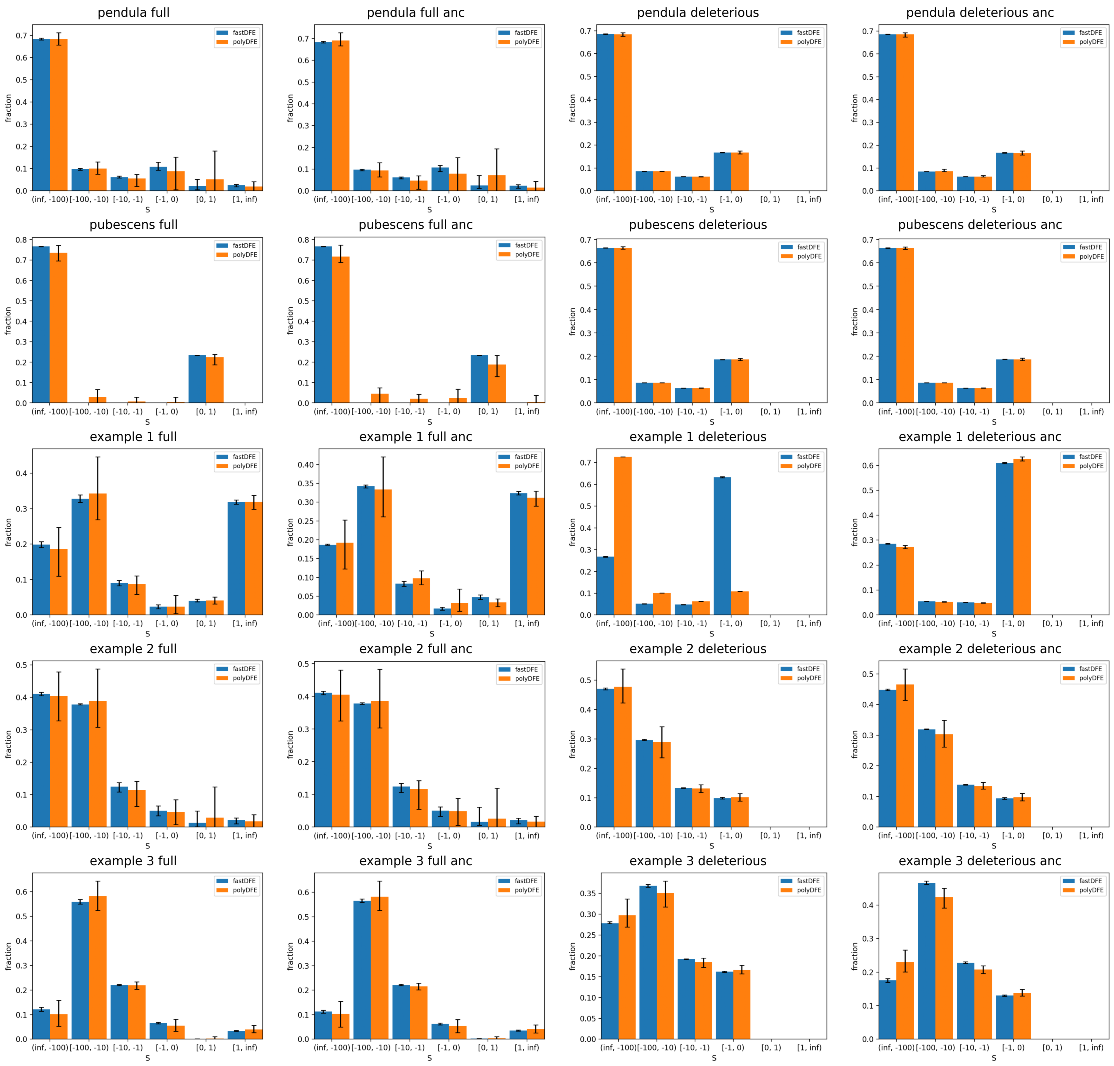


Figure B.1: Comparison of inferred discretized DFE between fastDFE and poldyDFE, employing the same datasets as in Table B.1. The underlying DFE parametrization is `GammaExpParametrization`. Each plot title indicates, in sequence, the sample set used, the DFE type inferred (beneficial or deleterious), and whether ancestral misidentification was included (*anc*). Vertical bars represent 95% confidence intervals. In most scenarios, the inferred DFE is very similar. One exception is `example 1 deleterious`, where polyDFE's optimization yields a likelihood of NaN, so that the initial values are reported. polyDFE's estimates also tend to have larger confidence intervals, likely due to less efficient numerical optimization.

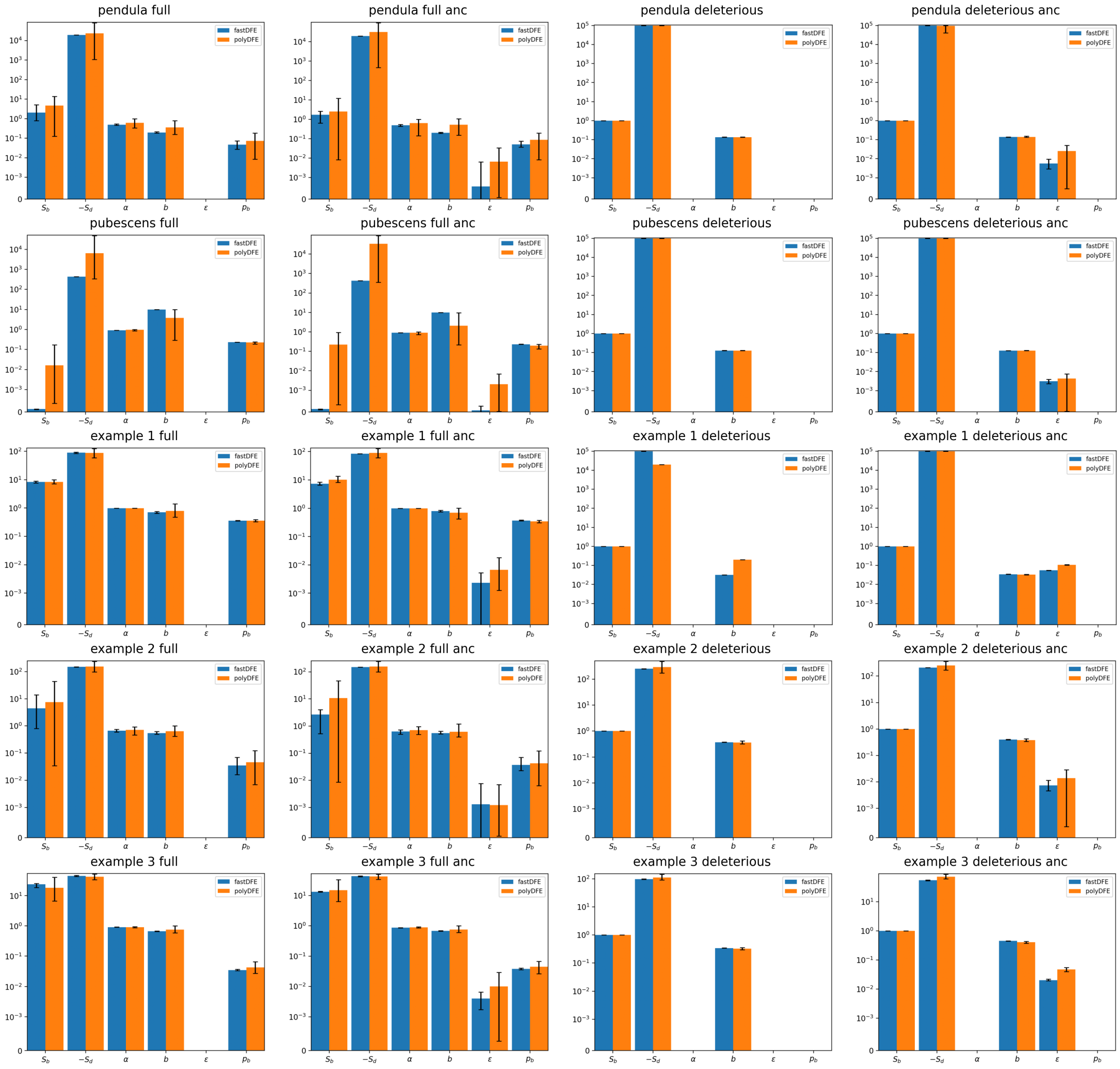


Figure B.2: The same comparison as in Figure B.1 but this time the inferred parameters are shown. Vertical bars represent 95% confidence intervals. Please note the logarithmic scale on the y-axis.

#### C Additional graphs

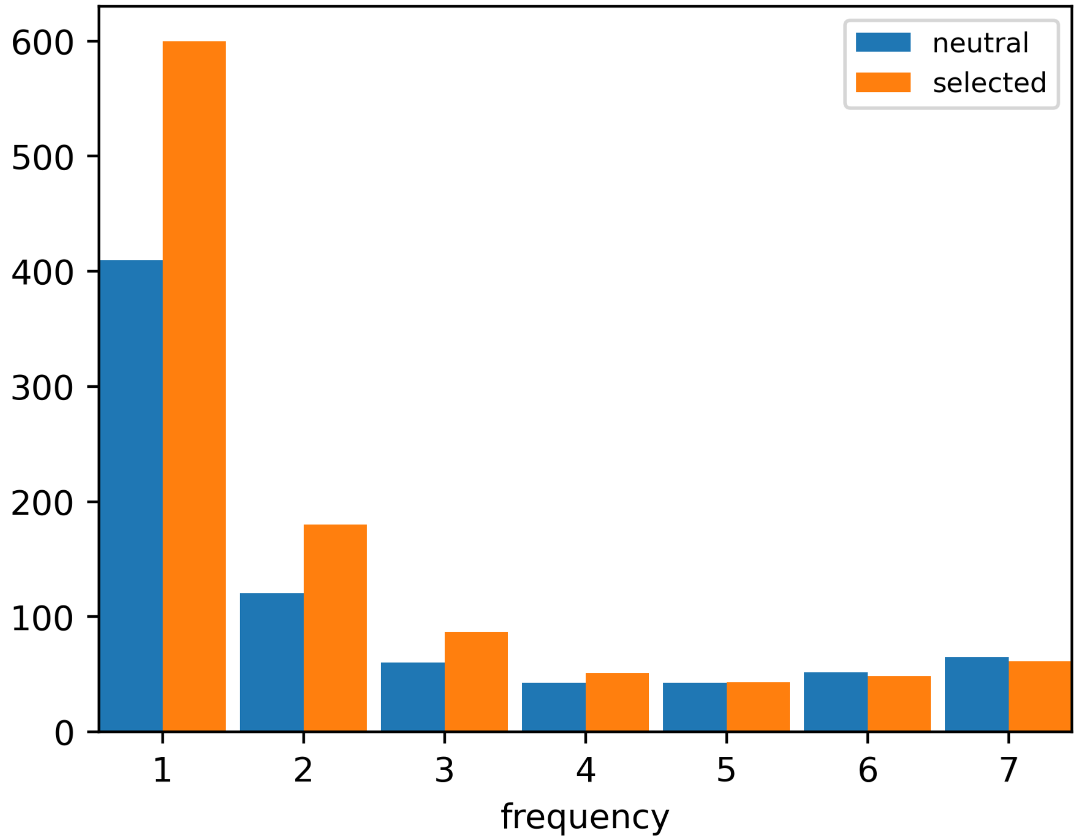


Figure C.1: SFS obtained from VCF file in code section in section 3.2.
